# Morning Elevation in Insulin Enhances Afternoon Hepatic Glucose Disposal in Dogs by Increasing Both Insulin Signaling and Glucose Action

**DOI:** 10.1101/2025.07.01.662587

**Authors:** Hannah L. Waterman, Marta S. Smith, Ben Farmer, Kalisha Yankey, Tristan Howard, Guillaume Kraft, Alan D. Cherrington, Dale S. Edgerton

## Abstract

The second-meal phenomenon refers to the improved glycemic response to a subsequent identical meal. We previously showed that morning (AM) hyperinsulinemia is a key mediator, priming the liver for enhanced net hepatic glucose uptake (NHGU) and glycogen storage during an afternoon (PM) hyperinsulinemic-hyperglycemic clamp. Postprandial NHGU is regulated by three primary mechanisms: insulin action (IA), initiated by hyperinsulinemia; glucose effectiveness (GE), driven by hyperglycemia; and the portal glucose signal (PGS), a neurally-mediated signal activated by glucose delivery into the hepatoportal circulation. It remained unclear, however, which of these mechanisms govern the increase in PM NHGU following AM insulin exposure. To address this, dogs underwent an AM clamp with either a 4-hour hyperinsulinemic prime (Prime, n=8) or basal insulin delivery (No Prime, n=8). After a 1.5-hour rest, both groups underwent a PM hyperglycemic clamp with portal glucose delivery under basal insulin conditions to isolate the effects of an AM insulin prime on PM glucose-mediated hepatic signals (GE/the PGS). Mean PM NHGU was significantly greater in the Prime group (2.2 ± 0.3 mg/kg/min) compared to the No Prime group (0.1 ± 0.3 mg/kg/min, p=0.005), accompanied by augmented net glycolytic and glycogen flux. These findings indicate that morning insulin can enhance glucose-mediated PM NHGU independently of a rise in PM insulin. However, maximal second-meal NHGU also requires elevated PM insulin. Together, this suggests that strategically timed early-day insulin or insulinotropic interventions could potentially improve hepatic responsiveness in settings of impaired postprandial glycemic control, such as insulin resistance or diabetes.

**Article Highlights:**

- Elevated morning insulin primes the liver for enhanced afternoon net hepatic glucose uptake (NHGU), but it was unclear whether augmentation of insulin action (IA), glucose effectiveness (GE), or the portal glucose signal (PGS) mediates this effect.
- Dogs underwent a morning euglycemic clamp with either elevated or basal insulin delivery, followed by an afternoon euinsulinemic-hyperglycemic clamp to isolate the effect of morning insulin priming on afternoon GE/PGS.
- Morning insulin priming enhanced afternoon NHGU via increased glucose-mediated mechanisms, though maximal afternoon NHGU also requires elevated afternoon insulin.
- These findings identify mechanisms underlying insulin-induced hepatic metabolic memory, providing a framework to inform strategies improving postprandial glucose handling in diabetes.

## Introduction

Effective postprandial glucose disposal is essential for maintaining metabolic health, and impairments in this process are a defining feature of insulin resistance and type 2 diabetes (1). Several factors influence postprandial hyperglycemia, including meal sequence and composition, rates of glucose disposal by the liver and peripheral tissues, glucose absorption from the gut, and rates of gastric emptying (2). These processes are integrated via the central nervous system and modulated by incretins, insulin, glucagon, and other blood glucose-regulating hormones (2).

An incompletely understood aspect of glucose regulation is the second meal phenomenon, in which the glycemic response to a second, identical meal is attenuated compared with its initial response to the same meal, suggesting that prior metabolic or hormonal cues influence subsequent glucose handling (3–5). Although this effect was first documented in humans in 1919 (6), the underlying mechanisms remain incompletely understood.

The liver is well positioned to contribute to this phenomenon, given its central role in postprandial glucose homeostasis. In response to nutrient ingestion, the liver both suppresses endogenous glucose production and accounts for a substantial portion of glucose disposal, with studies in humans and large animals indicating that the liver is responsible for the disposition of ∼60-65% of an oral glucose load when suppression of glucose production is considered (7–9). Thus, alterations in net hepatic glucose uptake (NHGU) and glycogen storage present a plausible mechanism linking prior meal exposure to subsequent glucose handling. Leveraging the experimental tools available in our laboratory to precisely quantify tissue-specific glucose uptake (7), we sought to identify regulators of this phenomenon.

Previously, we examined the effect of a morning (AM) duodenal glucose infusion, designed to mimic carbohydrates ingested at breakfast, on afternoon (PM) postprandial glucose metabolism, assessed with a hyperinsulinemic-hyperglycemic (HIHG) clamp (10). Dogs exposed to morning duodenal glucose exhibited a twofold increase in PM NHGU and hepatic glycogen deposition compared with saline controls, while non-hepatic glucose disposal was unaffected (10). To determine whether hepatic priming was driven by morning elevations in glucose, insulin, or gut-derived incretin signals, a follow-up study was performed to isolate those effects. Dogs underwent a morning pancreatic clamp in which either glucose or insulin were selectively elevated, mimicking what occurred during duodenal glucose infusion. All infusions were delivered into the hepatic portal vein to bypass the gut and incretin signaling. Morning hyperinsulinemia alone, independent of an AM rise in glucose and without incretin stimulation, nearly doubled NHGU during the PM clamp, recapitulating the response from our previous study (11). These findings indicate that morning hyperinsulinemia is a key mediator of the second meal phenomenon, priming the liver for enhanced glucose disposal later in the day by establishing hepatic metabolic memory.

NHGU is regulated by the integrated actions of three well-established primary signals: insulin action (IA), triggered by hyperinsulinemia; glucose effectiveness (GE), driven by hyperglycemia; and the portal glucose signal (PGS), activated by elevated glucose concentrations in the hepatoportal circulation relative to arterial circulation and related neural signaling, such as occurs with absorption of glucose from the gut after a meal (7; 11). Insulin promotes NHGU by suppressing endogenous glucose production and stimulating glucose uptake and glycogen synthesis through well-characterized signaling pathways (12). GE reflects the ability of glucose per se to suppress hepatic glucose output and enhance NHGU in the presence of basal insulin (13; 14). The PGS arises from afferent neural sensing of glucose delivery through the portal vein, enabling the liver to anticipate nutrient influx and adjust glucose uptake and storage accordingly, via neural mechanisms that are less well defined (7; 15). Together, these factors enable the liver to function as an active integrator of postprandial glucose metabolism (11).

Despite evidence that prior insulin exposure enhances NHGU later in the day, it remains unclear which afternoon regulatory signals (IA, GE, and/or the PGS) are augmented by morning insulin exposure. This distinction is particularly relevant because both insulin action and glucose action are impaired in insulin-resistance and type 2 diabetes, contributing to defective postprandial glucose disposal and exaggerated glycemic excursions in humans (16). Identifying the mechanisms underlying the priming effect of AM insulin is therefore important for defining how early hormonal cues regulate hepatic glucose metabolism, and may provide a framework for future strategies to potentially improve postprandial glucose handling and mitigate the impairments seen in metabolically dysregulated states. To address this knowledge gap, we assessed how an elevation in morning insulin influences glucose-mediated (GE and the PGS) versus insulin-mediated (IA) regulation of NHGU during a subsequent afternoon euinsulinemic-hyperglycemic clamp.

## Research Design and Methods

### Animal care and surgical procedures

16 adult mongrel dogs (8 males, 8 females; 27.6 ± 1.0 kg average body weight) were used. Animals were obtained from a USDA-approved vendor and housed according to American Association for the Accreditation of Laboratory Animal Care standards. All procedures were approved by the Vanderbilt Institutional Animal Care and Use Committee. Two weeks prior to experiments, dogs underwent a laparotomy under general anesthesia for the surgical placement of blood flow probes around the hepatic artery and hepatic portal vein. Sampling catheters were implanted for blood sampling in the hepatic vein, hepatic portal vein, and femoral artery. Infusion catheters were positioned in the inferior vena cava and the splenic and jejunal veins. All catheters were secured subcutaneously until the day of the experiment, at which time they were exteriorized under local anesthesia. The dogs were fed a controlled diet consisting of chow and meat (46% carbohydrates, 34% protein, 14.5% fat, and 5.5% fiber) and fasted for 18h prior to study. Assessments were performed the day before the experiment to confirm health, requiring ≥75% meal consumption, leukocytes <18,000/mm^3^, and hematocrit >34%. Blood collection was limited to ≤20% of the animal’s total blood volume.

### Experimental Design

The experimental protocol lasted 8h in total, consisting of a 4h morning clamp period (0-240 min) followed by a 90-min break (240-330 min) and a subsequent 2.5h afternoon clamp period (330-480 min; **Fig. 1).** Blood samples were collected every 15-30 min from the femoral artery, hepatic portal vein, and the left major hepatic vein for measurement of hormone and metabolite levels. Arterial plasma glucose was assessed every 5 min, and peripheral leg vein glucose infusion rates were adjusted as needed to maintain target plasma glucose concentrations.

**Figure 1:**
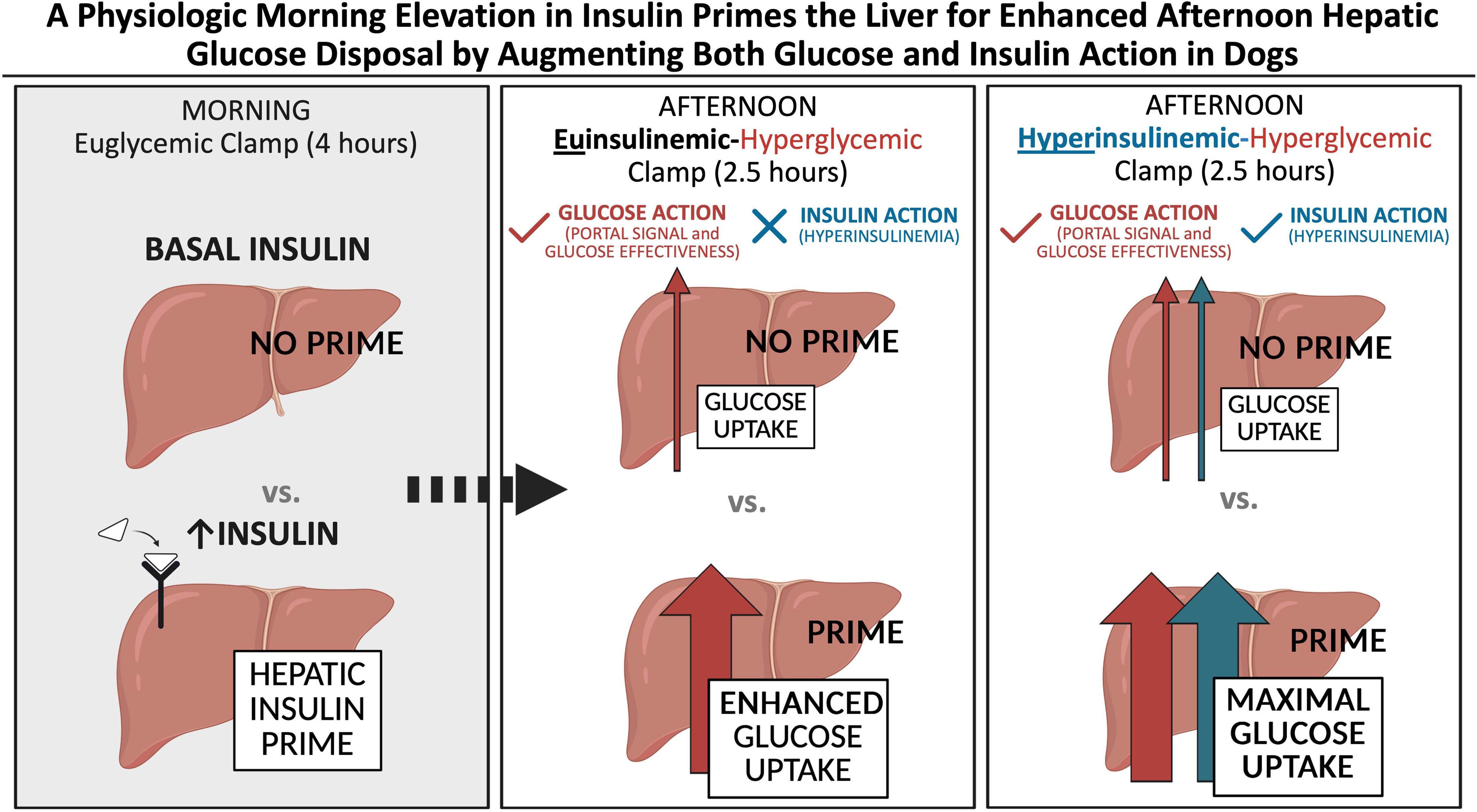
Experimental protocol overview. Canines (*n*=8/group) underwent a 4h euglycemic clamp in the morning (0 to 240 min) with either basal insulin (No Prime; orange) or hyperinsulinemia (Prime; green). Near the end of the AM clamp (180 min), tracer infusion began to allow enough time for equilibration before the start of the PM clamp. Following the AM clamp, there was a 1.5h rest period (240-330 min) where all infusions except tracer were halted. The dogs then underwent a 2.5h PM hyperglycemic clamp with portal glucose delivery and basal insulin (330 to 480 min). Hepatic tissue collection occurred at the end of the PM clamp. IV; intravenous infusion.

### Morning (AM) Clamp Period (0-240 min) and Non-Clamp Period (240-330 min)

To suppress endogenous pancreatic insulin and glucagon secretion, somatostatin (0.8 µg/kg/min; Bachem, Torrance, CA) was infused into the inferior vena cava at the onset of the AM clamp. Glucagon (0.57 ng/kg/min; GlucaGen, Boehringer Ingelheim, Ridgefield, CT) was replaced intraportally to maintain basal plasma levels. The Prime group received an elevated intraportal insulin infusion (Novolin R; Novo Nordisk, Basværd, Denmark) designed to replicate physiologic secretion observed after a morning duodenal glucose infusion previously observed (2.1 mU/kg/min from 0-30 min, 2.4 mU/kg/min from 30-60 min, and 1.5 mU/kg/min from 60-240 min) (17). The No Prime group received basal intraportal infusion replacement (0.25 mU/kg/min). Prior research from our lab has shown that the enhancement of PM NHGU is driven by an increase in AM insulin levels, rather than a rise in morning glucose levels (17). Therefore, arterial plasma glucose was clamped at euglycemia (∼110 mg/dL) for the entirety of the AM clamp. [3-^3^H] glucose (Revvitty, Waltham, MA) was infused peripherally starting at 180 min in both groups (38 µCi priming, 0.38 µCi/min continuous) to enable tracer-based assessment of direct hepatic glycogen synthesis.

After the AM clamp (240 min), all infusions stopped for 1.5h except tracer. Blood samples were obtained from the femoral artery, portal vein, and hepatic vein at 300, 315, and 330 min to define the baseline hepatic metabolic state prior to the PM clamp **(Fig. 1).**

### Afternoon (PM) Clamp Period (330-480 min)

Both groups underwent an identical afternoon 2.5h hyperglycemic clamp with basal replacement of insulin and glucagon **(Fig. 1).** Somatostatin (0.8 µg/kg/min) was infused via a peripheral vein, while basal glucagon (0.57 ng/kg/min) and basal insulin (0.25 mU/kg/min) were delivered intraportally. To activate the portal glucose signal, glucose was infused directly into the portal vein (4 mg/kg/min). A primed, continuous peripheral glucose infusion quickly raised plasma glucose levels to ∼230 mg/dL, which was maintained for the duration of the PM clamp. All dogs were anesthetized with infusions still ongoing at the end of the PM clamp (480 min) to allow for accurate tissue sampling reflective of the PM clamp conditions. Hepatic tissue was rapidly excised, flash-frozen in liquid nitrogen, and stored at -80°C for subsequent molecular analyses. The animals were then euthanized.

### Analyses

#### Biochemical and Molecular Methods

Whole blood was used for arteriovenous balance measurements (18). Plasma glucose was measured in quadruplicate using an Analox GM9 analyzer. Plasma insulin (#PI-12K, MilliporeSigma, Burlington, MA), glucagon (#GL-32K, MilliporeSigma), and cortisol (MP Biomedicals, Santa Ana, CA) were measured by radioimmunoassay (VUMC Analytical Services Core) (18). Glucagon values were background-corrected for non-specific binding.

Metabolites (lactate, glycerol, alanine, and non-esterified fatty acids) were measured using enzymatic spectrophotometric assays as previously described (18). Plasma [3-^3^H]-glucose specific activity was determined after deproteinization using the Somogyi method and quantified by liquid scintillation counting (19). Terminal hepatic glycogen content was measured using the Keppler-Decker amyloglucosidase method (20).

Terminal liver tissue was analyzed by qPCR, western blotting, and glucokinase activity assays. Overnight-fasted dogs (n=5) served as basal references. Gene expression was normalized to glyceraldehyde 3-phosphate dehydrogenase (GAPDH), and protein levels were normalized to total protein using Ponceau S staining (18). All assays were optimized for canine-specific targets using validated primers and antibodies (18). Glucokinase activity was measured using an assay adapted from Shiota et al. (21). Detailed protocols, including reagents, primers, antibodies, sample preparation, and calculations, are provided in the Supplemental Materials.

#### Calculations

Arterio-venous difference calculations were used to determine net hepatic glucose balance (NHGB) where NHGB = [HGLout] - [HGLin]. Glucose output from the liver (HGLout) was calculated as [HGLout] = [BFh x Gh], while glucose entering the liver (hepatic sinusoidal) was calculated as [HGLin] = [(BFa × Ga) + (BFp × Gp)]. Here, G denotes blood glucose concentration; BF denotes blood flow; A, P, and H correspond to the hepatic artery, hepatic portal vein, and hepatic vein, respectively. Plasma values were converted to whole blood using a factor of 0.73 (18). Negative NHGB indicates net hepatic glucose uptake (HGLin>HGLout), as reported here. This approach was also used to assess net hepatic balance for all other substrates and hormones. Hepatic fractional glucose extraction was calculated as NHGB/HGLin. Non-hepatic glucose uptake was derived as GIR – NHGU, corrected for changes in glucose mass. Direct hepatic glycogen synthesis was calculated as radiolabeled glycogen content divided by specific activity of the glucose precursor pool during the PM clamp. The specific activity of the glucose precursor pool was determined from the ratio of labeled to unlabeled glucose in inflowing blood, weighted by hepatic artery and portal vein contributions.

Net glycogen flux (net glycogenesis/glycogenolysis) was calculated as NHGU + NHLU + 2(NHAU) + NHGlyU – HGO where NHLU, NHAU, and NHGlyU represent net hepatic uptake of lactate, alanine, and glycerol, respectively, and HGO represents hepatic glucose oxidation. In conditions of net hepatic lactate output (NHLO), NHLU becomes negative. NHAU was doubled to account for total amino acid uptake, as alanine accounts for ∼50% of gluconeogenic amino acid flux (22). HGO was assumed to be ∼0.2 mg/kg/min under these conditions, contributing minimally to overall flux (23). Net glycolytic/gluconeogenic flux was calculated as glycolytic flux – gluconeogenic flux, where glycolytic flux = NHLO + 0.1(NHLO) + HGO and gluconeogenic flux = 2(NHAU) + NHGlyU. Pyruvate flux was approximated as 10% of lactate flux (24).

#### Statistics

Data are expressed as mean ± SEM. Group differences and temporal changes in flux were analyzed using a two-way repeated-measures ANOVA with Tukey’s post hoc test for multiple comparisons. Area under the curve (AUC) was compared using an unpaired two-tailed t-test, and molecular outcomes were analyzed using a one-way ANOVA with Tukey’s post hoc correction. Normality of all datasets was confirmed using the Shapiro-Wilk test, and all data were found to follow a normal distribution. A threshold of p < 0.05 was used to define statistical significance. All analyses were performed using GraphPad Prism software.

#### Data and Resource Availability

All data points are reported in the Supporting Data Values file. Data and supplemental materials are publicly available at 10.6084/m9.figshare.31049578. Additional data are available from the corresponding author upon request.

## Results

### AM Clamp Glucose and Hormone Data

During the AM clamp, euglycemia (105-115 mg/dL) was maintained in both groups **(Fig. 2A).** Arterial plasma insulin was kept at basal levels in the No Prime group (6 ± 1 µU/mL), whereas insulin was elevated in the Prime group according to the experimental design (32 ± 3 µU/mL, p<0.0001; **Fig. 2B, C**; **Table 1**). Accordingly, hepatic sinusoidal insulin concentrations were approximately sixfold higher in the Prime group than in the No Prime group (**Fig. 2D, E**; **Table 1**). Arterial and hepatic sinusoidal glucagon concentrations remained basal and did not differ between groups **(Table 1).** Consistent with the higher exposure to insulin, the Prime group required significantly more exogenous glucose to maintain euglycemia during the AM clamp than the No Prime group **(Fig. 2F, G**; **Table 1).** Of note, AM net gut glucose uptake was not different between the two groups (0.3 ± 0.1 vs. 0.2 ± 0.1 mg/kg/min in Prime vs. No Prime, respectively). Thus, the AM clamp successfully established sustained hyperinsulinemia in the Prime group compared to the No Prime group without altering circulating glucose or glucagon levels.

**Figure 2:**
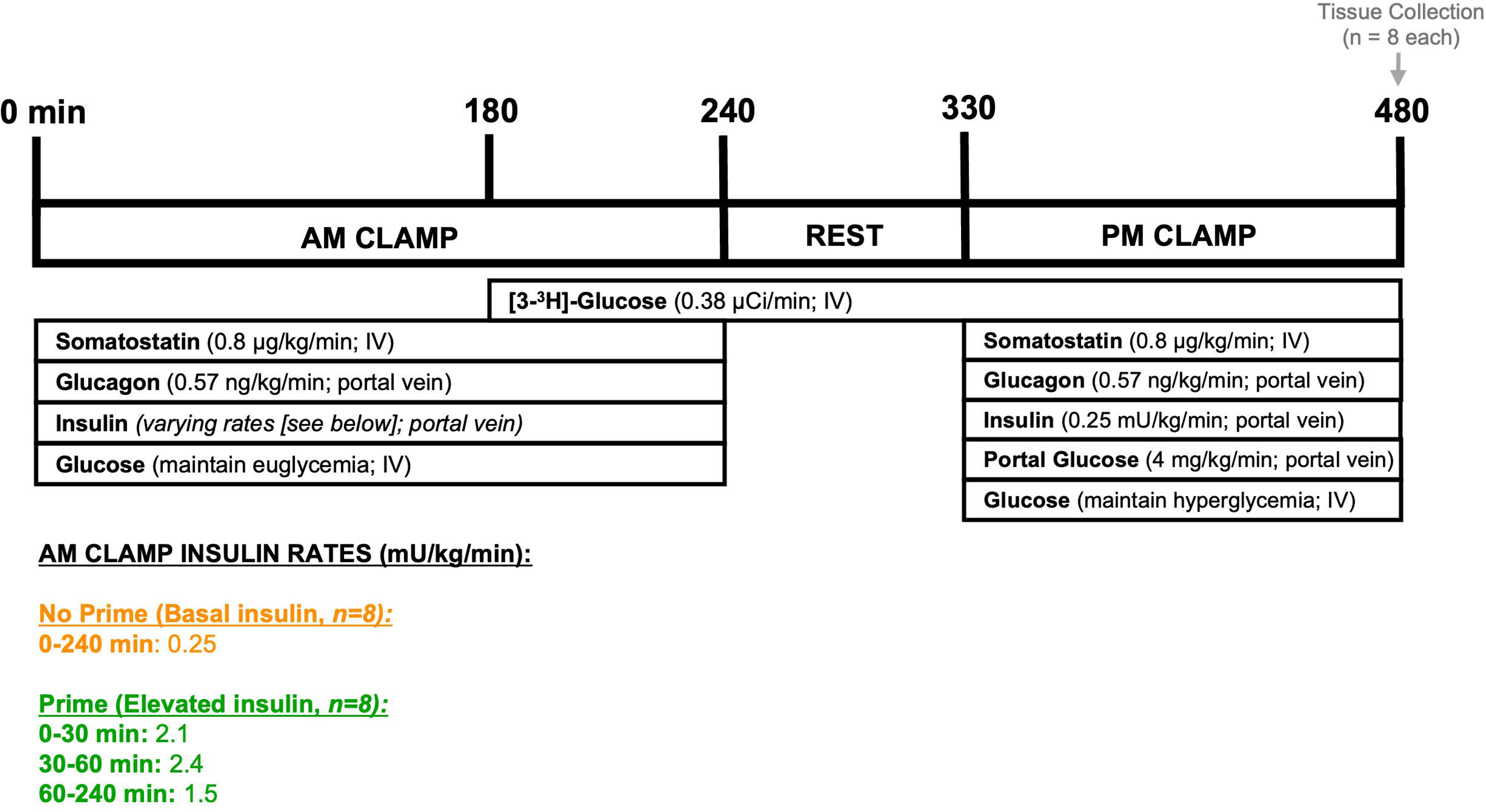
Morning (AM) clamp glucose and insulin flux data. Arterial plasma glucose (A) and insulin (B), hepatic sinusoidal plasma insulin (D), and the amount of exogenous glucose required to maintain euglycemia (F) are shown for the No Prime and Prime groups; *n*=8/group. Bar graphs indicate the area under the curve (AUC) for arterial insulin concentrations (C), hepatic sinusoidal insulin concentrations (E), and glucose infusion rates (G) during the 4h AM period. Data are expressed as mean ± SEM. ****P<0.0001 between groups. ns = non-significant. No prime; orange. Prime; green.

**Table 1.**
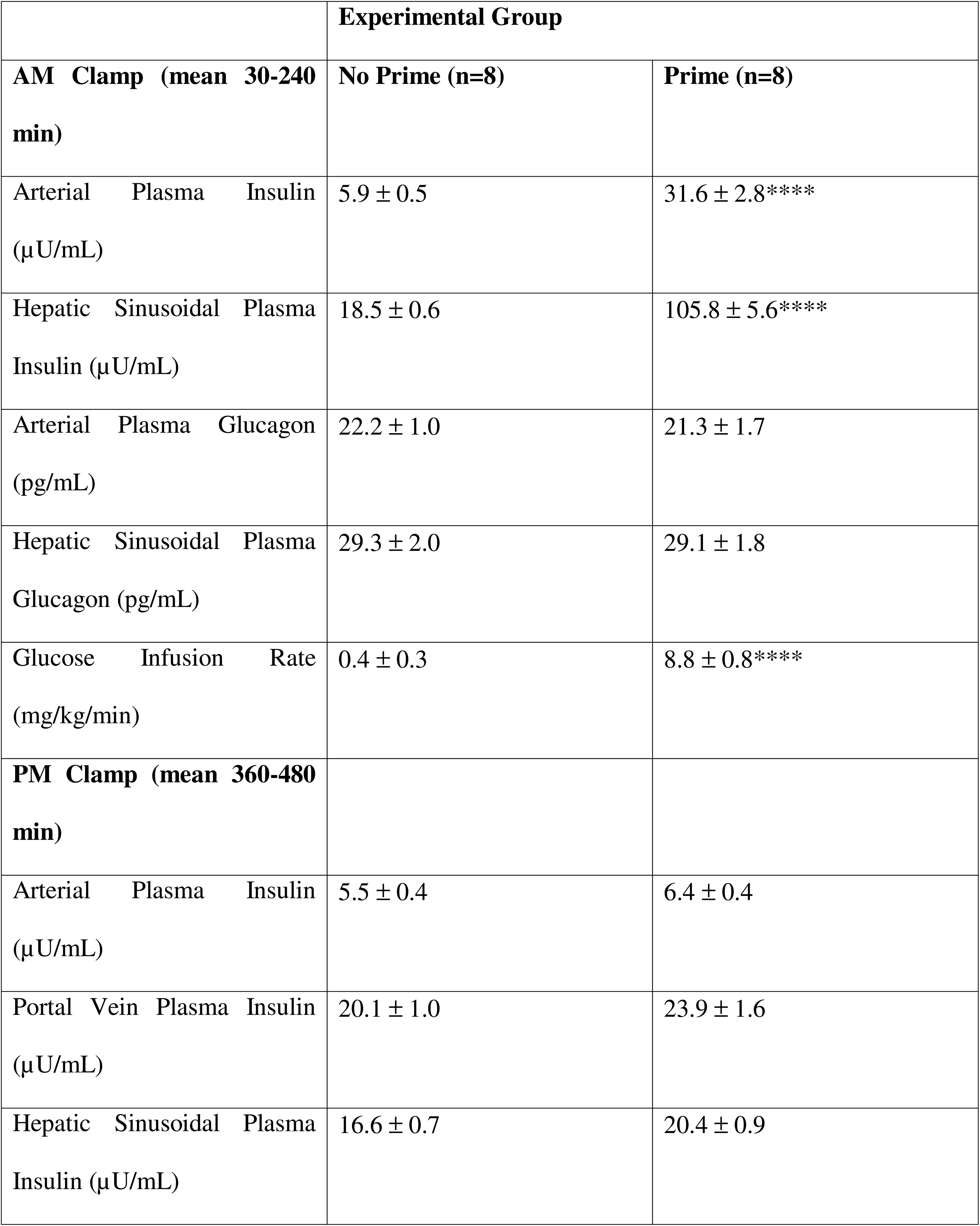
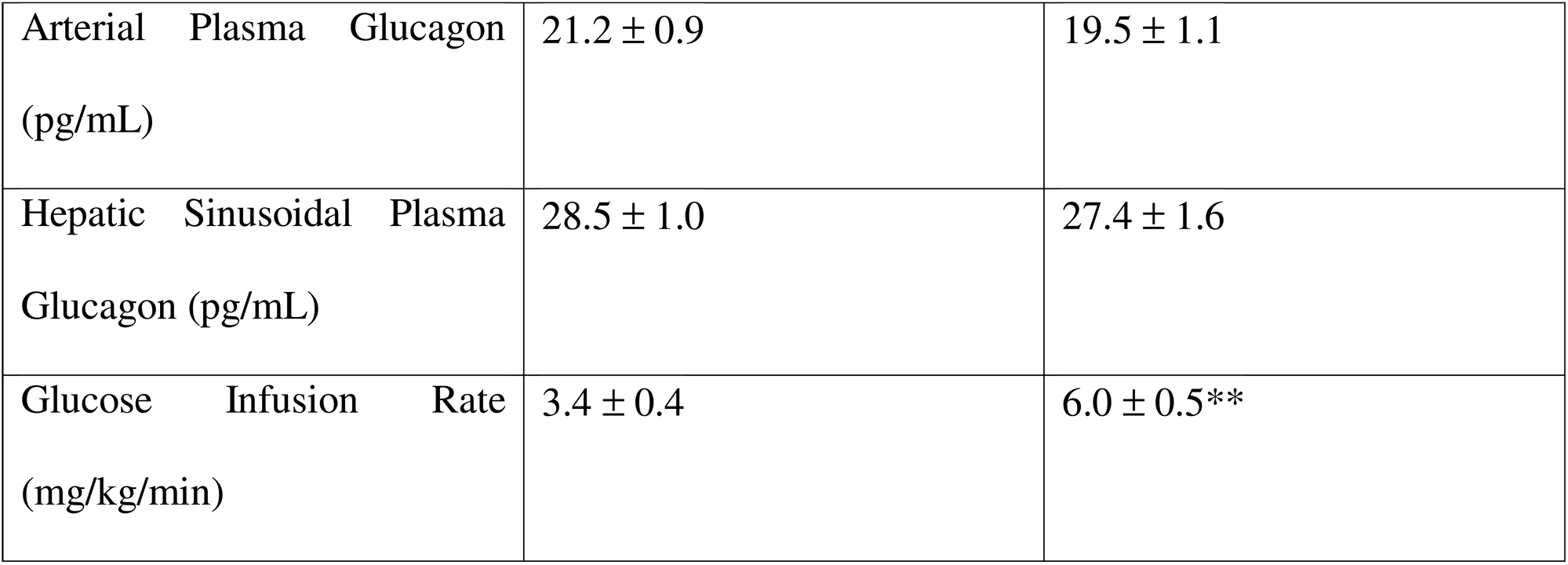
Mean plasma insulin and glucagon levels in the arterial and hepatic sinusoidal circulation and glucose infusion rate during the AM and PM clamps.

### PM Clamp Glucose, Hormone, and Metabolite Data

During the PM clamp, experimental conditions were designed to be matched between groups. Arterial plasma glucose was clamped at a hyperglycemic level (230-240 mg/dL; **Fig. 3A, B**), the portal glucose signal was activated by delivering glucose intraportally, resulting in a negative arterial-to-portal glucose gradient **(Fig. 3C)**, and arterial and hepatic sinusoidal insulin concentrations were maintained at basal levels in both groups **(Fig. 3D**; **Table 1).** Despite similar PM clamp conditions, the Prime group required 76% more glucose to maintain the hyperglycemic clamp than the No Prime group (glucose infusion rate of 6.0 ± 0.5 vs. 3.4 ± 0.4 mg/kg/min, respectively, p=0.006; **Fig. 4A, B**; **Table 1).** This difference was not attributable to changes in non-hepatic glucose uptake, which did not differ between groups (3.6 ± 0.3 vs. 3.1 ± 0.3 mg/kg/min in Prime vs. No Prime, respectively; **Fig. 4C, D).** Net glucose uptake by the gut was not different between the two groups (0.5 ± 0.3 mg/kg/min in Prime vs. 0.4 ± 0.1 mg/kg/min in No Prime), either. Instead, the increased glucose requirement was due to a marked increase in net hepatic glucose uptake (NHGU) in the Prime group (mean of 2.2 ± 0.3 vs. 0.1 ± 0.3 mg/kg/min in Prime vs. No Prime, p=0.005; **Fig. 4E, F).** The enhancement in PM NHGU in the Prime group reflected both increased net glycolytic flux **(Fig. 4G)** and increased net glycogenesis **(Fig. 4H).** Thus, morning insulin priming resulted in a sustained amplification of PM hepatic glucose uptake even in the absence of a rise in PM insulin levels.

**Figure 3:**
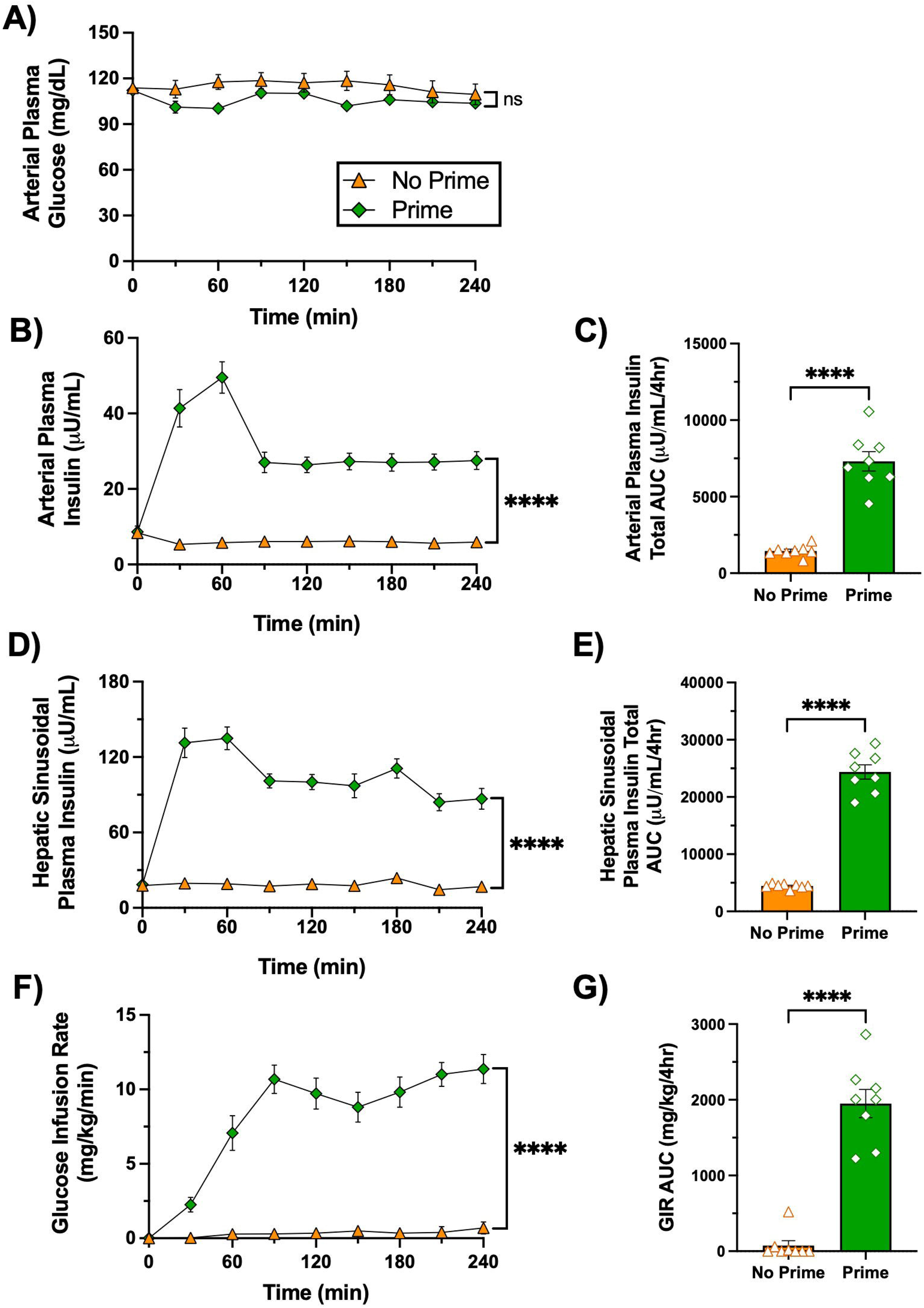
Afternoon (PM) clamp glucose and insulin data. A vertical line at 330 min separates the rest period from the onset of the PM clamp. Arterial plasma glucose (A), hepatic glucose load (B), the difference between the arterial and the hepatic portal vein plasma glucose levels (C), and hepatic sinusoidal plasma insulin (D) are shown for the 2.5h euinsulinemic-hyperglycemic PM clamp period in the No Prime and Prime groups; *n*=8/group. There were no statistically significant differences between the two groups for any of the parameters measured. Data are expressed as mean ± SEM. ns = non-significant. No prime; orange. Prime; green.

**Figure 4:**
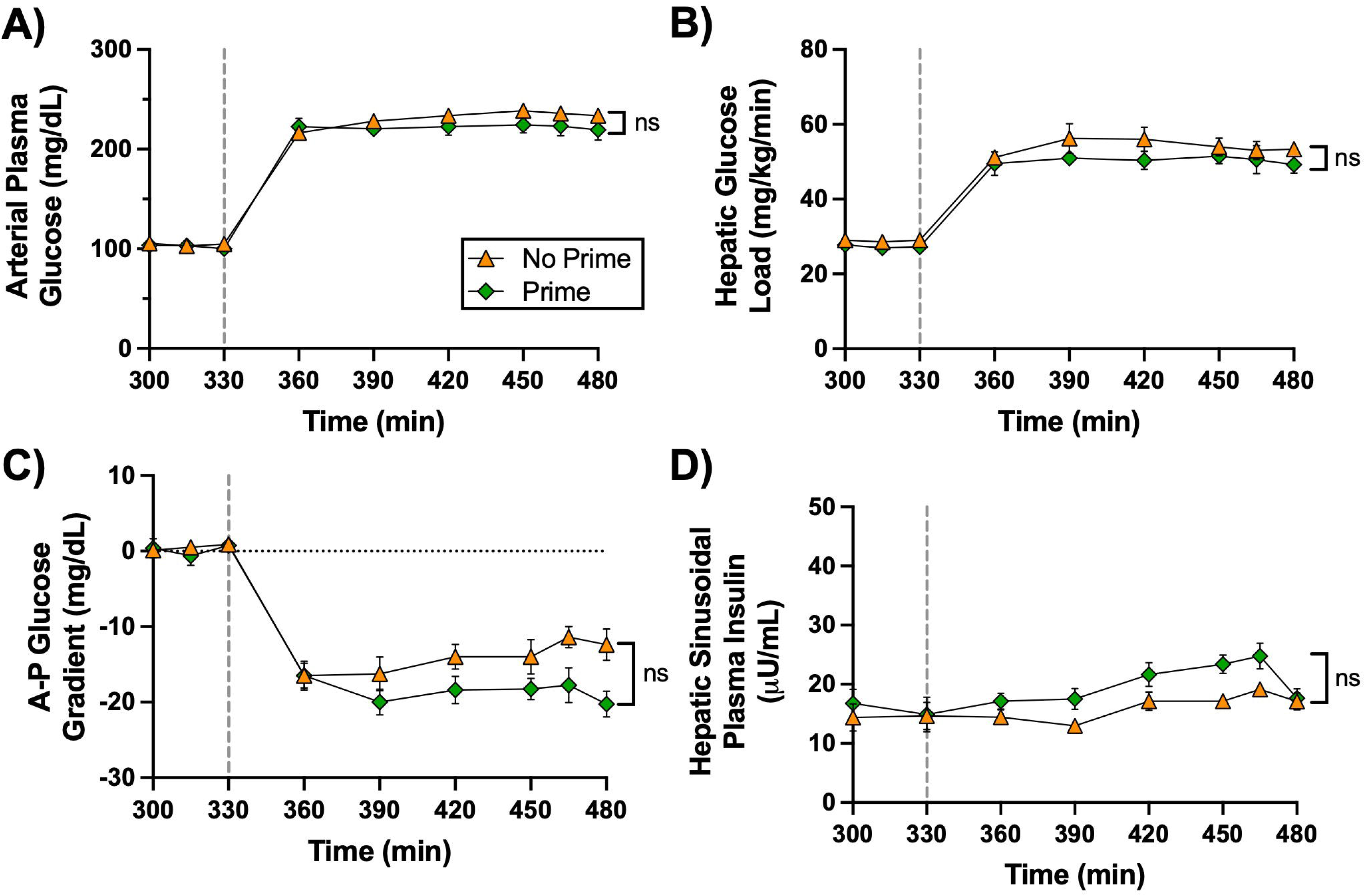
Glucose uptake and storage during the PM euinsulinemic-hyperglycemic clamp. A vertical line at 330 min separates the rest period from the onset of the PM clamp. Glucose infusion rate (A), non-hepatic glucose uptake (C), net hepatic glucose uptake (E), net glycolytic/gluconeogenic flux (G), and net glycogenesis/glycogenolysis flux (H) are shown over time for the No Prime and Prime groups, *n*=8/group. The respective delta AUCs for the glucose infusion rate, non-hepatic glucose uptake, and net hepatic glucose uptake are shown for both groups (B, D, F). Data are expressed as mean ± SEM. *P<0.05, **P<0.01, ***P<0.001 between groups. ns = non-significant. No prime; orange. Prime; green.

Arterial non-esterified fatty acid (NEFA) concentrations were significantly higher in the No Prime group than in the Prime group during the PM clamp, whereas net hepatic NEFA uptake did not differ between groups **(Fig. 5A, B).** Arterial glycerol concentrations and net hepatic glycerol uptake followed a similar trend but were not significantly different between groups **(Fig. 5C, D).** Arterial alanine concentrations were comparable; however, net hepatic alanine uptake was reduced modestly in the Prime group relative to the No Prime group **(Fig. 5E, F).** Arterial lactate concentrations did not differ between groups, but net hepatic lactate output was approximately fivefold greater in the Prime group than in the No Prime group **(Fig. 5G, H),** consistent with increased hepatic glucose uptake and glycolytic flux. For many of these parameters, the Prime and No Prime groups did not begin at the same baseline level at 330 min.

**Figure 5:**
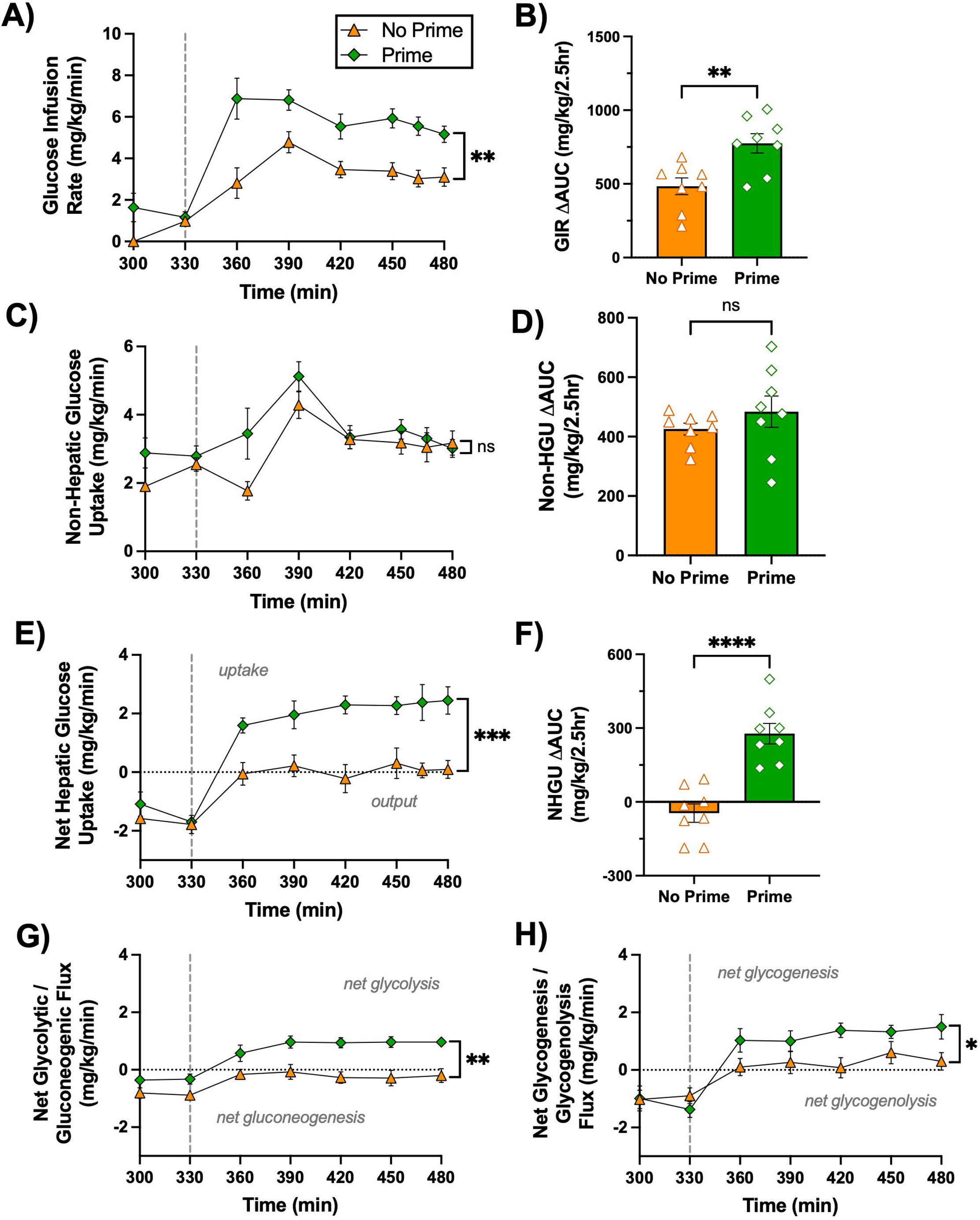
PM clamp fatty acid and metabolite flux data. A vertical line at 330 min separates the rest period from the onset of the PM clamp. Arterial plasma non-esterified fatty acids (NEFA) (A), net hepatic NEFA uptake (B), arterial blood glycerol (C), net hepatic glycerol uptake (D), arterial blood alanine (E), net hepatic alanine uptake (F), arterial blood lactate (G), and net hepatic lactate output (H) are shown for the PM euinsulinemic-hyperglycemic clamp for the No Prime and Prime groups, *n*=8/group. Data are expressed as mean ± SEM. **P<0.01 between groups. ns = non-significant. No prime; orange. Prime; green.

### Hepatic Glycogen Content, Transcript, and Protein Expression

Terminal hepatic glycogen content was significantly greater in the Prime group compared with the No Prime group (43 ± 3 vs. 36 ± 1 mg glycogen/g liver, p=0.014; **Fig. 6A**). However, direct glycogen synthesis during the PM clamp was not statistically significantly different between groups **(Fig. 6B).** Phosphorylated Akt relative to total Akt was similar in both the Prime and No Prime groups when compared to overnight-fasted basal dogs, consistent with basal insulin conditions during the PM clamp **(Fig. 6C).** *Glucokinase* (*GCK*) mRNA abundance was significantly greater in the Prime group compared to both the No Prime and basal groups at the end of the PM clamp **(Fig. 6D);** however, GCK protein abundance and enzymatic activity did not differ between any of the groups **(Fig. 6E, F).** Glycogen synthase protein was similarly dephosphorylated and activated in both experimental groups relative to basal dogs **(Fig. 6G),** while phosphorylated glycogen phosphorylase protein did not differ among groups **(Fig. 6H).** Transcript levels of *phosphoenolpyruvate carboxykinase 1* (*PCK1*) and *glucose-6-phosphatase catalytic subunit 1* (*G6PC1*) were similarly reduced in both Prime and No Prime groups compared with basal **(Supplemental Fig. 1A, 1B).** In contrast, *sterol regulatory element-binding protein-1c (SREBP-1c)* and *fatty acid synthase (FAS)* mRNA levels were significantly elevated in the Prime group relative to the No Prime group at the end of the PM clamp **(Supplemental Fig. 1C, 1D).** Respective western blot images can be viewed in **Supplemental Fig. 1E**.

**Figure 6:**
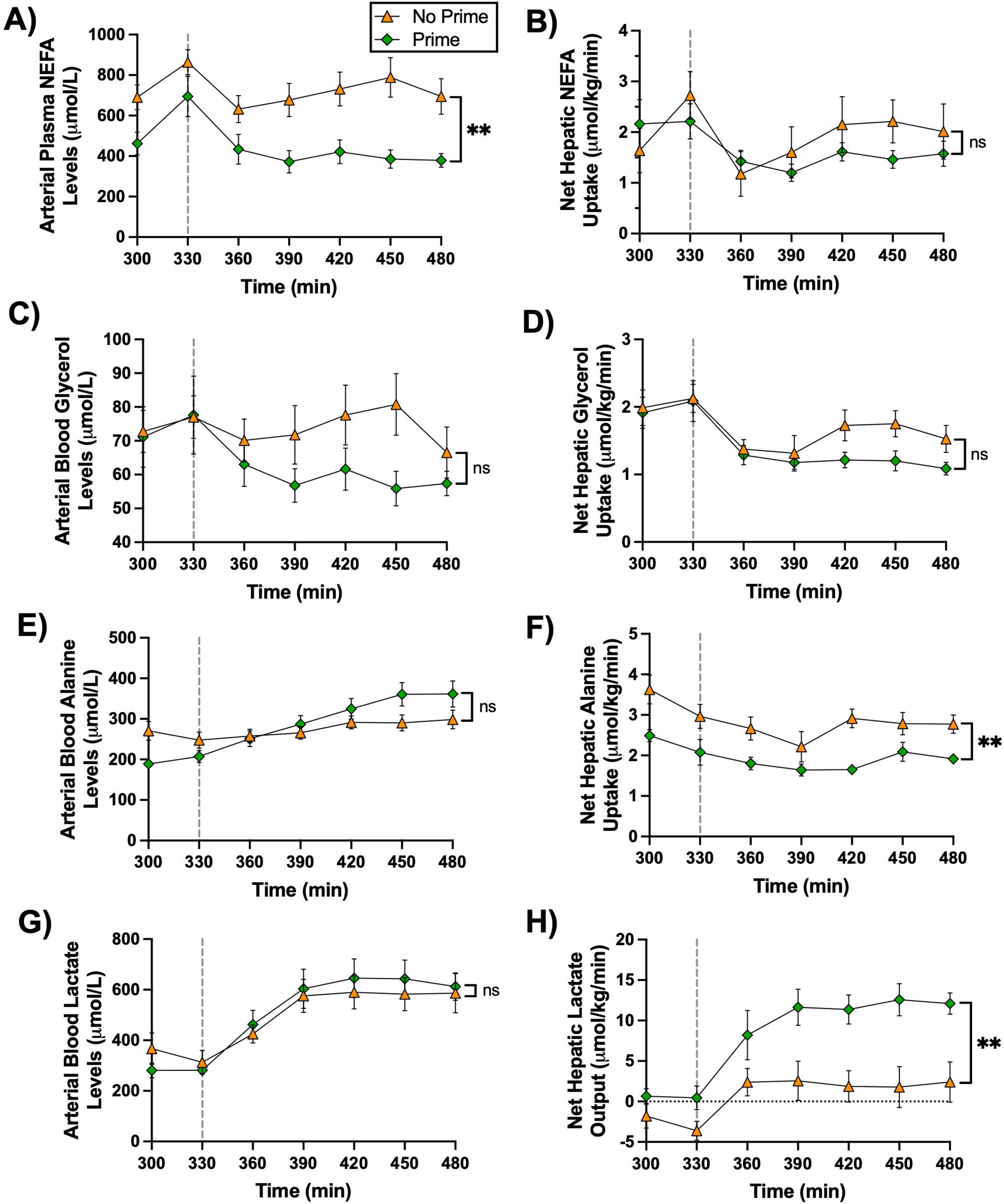
Glycogen content and hepatic gene/protein expression at the end of the PM clamp. Terminal hepatic glycogen content is shown in (A), and tracer-determined direct glycogen synthesis during the PM clamp is shown in (B). Protein and gene expression are shown as follows: Akt protein (phospho/total) in (C), glucokinase (GCK) mRNA in (D), GCK protein (normalized to Ponceau S stain) in (E), glucokinase (GCK) activity at 0.5 Vmax in (F; n = 5 or 6/group), glycogen synthase (GS) protein (phospho/total) in (G), and glycogen phosphorylase (GP) protein (phospho/total) in (H). Protein data were obtained by western blot and normalized to Ponceau S stain; representative images are provided in Supplemental Figure 1. Data are presented for the No Prime and Prime groups (n = 8/group), with basal tissue values included where applicable (n = 5). All values are expressed as mean ± SEM. *P < 0.05, **P < 0.01, ***P < 0.001, ****P < 0.0001 between groups; ns = not significant. Basal; grey, No prime; orange. Prime; green.

## Discussion

The second meal phenomenon, in which the glycemic response to a subsequent identical meal is attenuated compared with the first, is a mechanism by which the body integrates prior hormonal and nutrient exposures to optimize later postprandial glucose handling (4; 5; 25). First observed in humans in the early 20th century, sequential glucose tolerance tests revealed progressively smaller glycemic excursions with repeated carbohydrate loads of the same size (6). Over the decades, multiple mechanisms have been proposed, including enhanced insulin secretion during the second meal, gut-derived incretin effects, improved skeletal muscle insulin sensitivity, and increased muscle glycogen content (4; 26-29). Liver glycogen accumulation has also been suggested to play a key role, as sequential glucose loads appear to progressively augment hepatic carbohydrate storage (4; 30). Despite these observations, human studies have largely relied on indirect or correlative measures, such as circulating hormone levels. As a result, the relative roles of hepatic versus extrahepatic tissues, and the contributions of hormonal versus nutrient-driven priming, remained unclear. Using a conscious dog model, which allows precise control of circulating glucose and insulin via pancreatic clamps while bypassing incretin signals, we previously showed that a physiological morning insulin rise primes the liver for enhanced NHGU and glycogen storage during a later hyperglycemic clamp, without affecting non-hepatic glucose uptake (17; 31). These findings support insulin-driven hepatic metabolic memory as a key contributor to the second-meal phenomenon and suggest that interventions targeting the timing or amplitude of insulin exposure could improve postprandial glucose handling, particularly in diabetes (3; 5; 25; 32-34).

To dissect the physiological mechanisms of morning insulin priming, we used a reductive approach to distinguish the contributions of glucose- versus insulin-mediated signals. For context, prior work (10; 35) in which all three regulatory factors (IA, GE, and the PGS) were engaged under controlled postprandial conditions provides a reference for the full priming effect, where morning insulin priming increased PM NHGU by 3.8 mg/kg/min **(Fig. 7A).** In the present study, with matching hyperglycemic levels and portal glucose delivery but under basal insulin conditions, morning priming increased PM NHGU by 2.1 mg/kg/min (reflecting the effect of AM insulin on GE and/or the PGS; **Fig. 7B).** These results indicate that approximately half of the priming effect is mediated by enhanced responsiveness to glucose-mediated signals, while the remaining portion reflects augmented afternoon insulin action (1.7 mg/kg/min) **(Fig. 7C).** In the No Prime group, when suppression of hepatic glucose production is considered, glucose- and insulin-mediated signals contribute similarly to PM NHGU. Thus, morning insulin priming appears to proportionally enhance both pathways rather than altering their relative contributions to PM NHGU. Although this calculation relies on cross-study comparisons and assumes additive effects, it highlights that morning insulin exposure amplifies the liver’s sensitivity to both hormonal and glucose cues later in the day, enhancing postprandial glucose disposal (36). Overall, these findings demonstrate that morning insulin augments both glucose-mediated (GE and/or PGS) and insulin-mediated (IA) components of afternoon NHGU.

**Figure 7:**
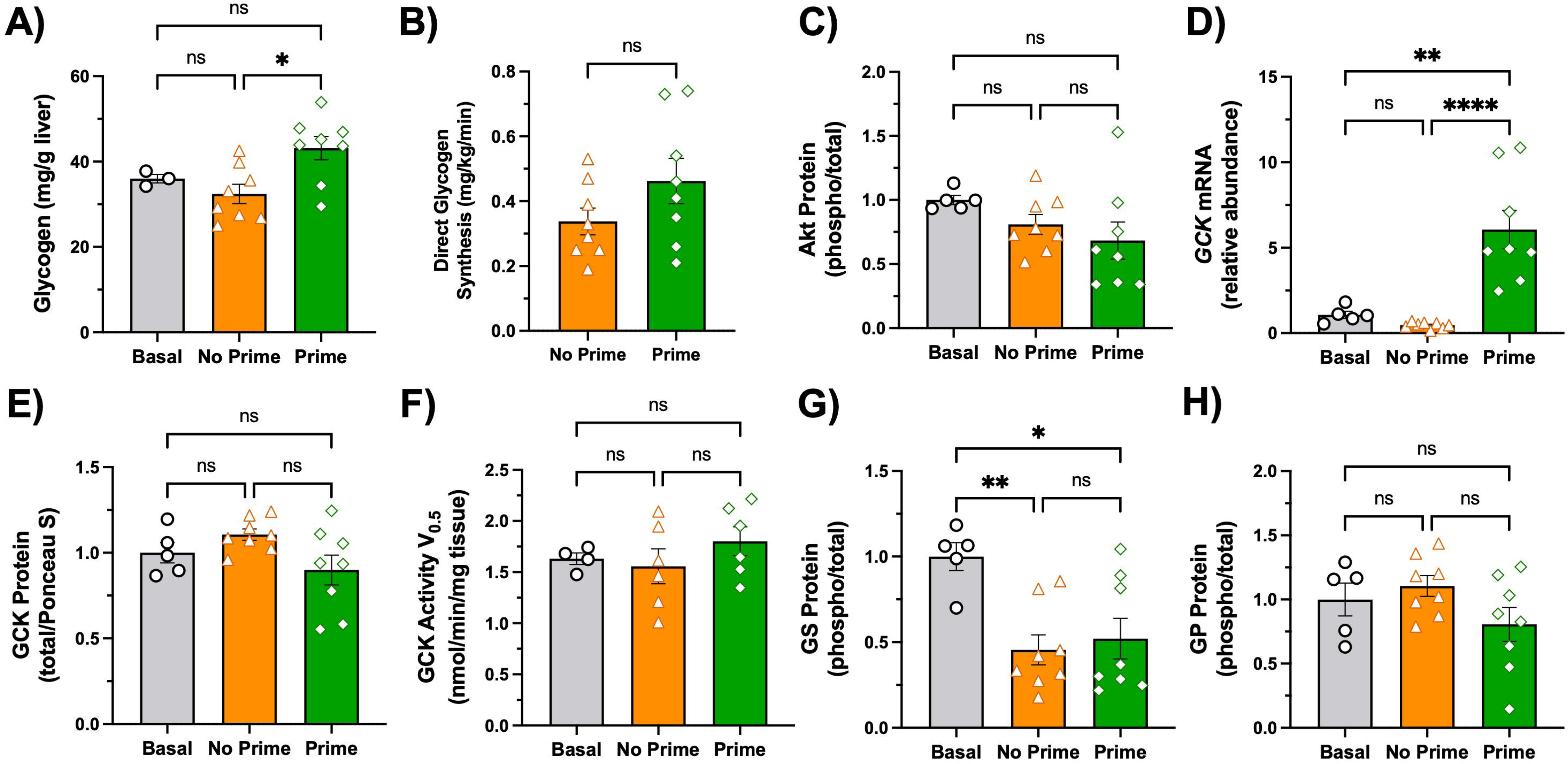
Summary of PM net hepatic glucose uptake (NHGU) data. Mean PM NHGU data from control experiments previously completed are shown in (A). One of these groups (grey) received basal insulin in the AM with a PM HIHG clamp. The other (blue) received hyperinsulinemia in the AM with the same PM HIHG clamp. The size of the priming effect of AM insulin on PM NHGU is the difference in PM NHGU between

Although the PM clamp conditions were designed to be matched between groups, random, non-significant differences were observed in the arterial-portal (A-P) glucose gradient and hepatic sinusoidal insulin levels. In the current study, the PM A-P glucose gradient was 5 mg/dL smaller and hepatic sinusoidal insulin levels were 3.8 µU/mL lower in the No Prime group compared with the Prime group. Prior studies using similar hyperglycemic clamps with portal glucose infusion have shown that NHGU scales directly with hepatic glucose load and responds linearly to rises in hepatic insulin within the physiological range (36; 37). Based on these data, the observed differences would be expected to result in at most a ∼0.5 mg/kg/min underestimation of PM NHGU in the No Prime group. These minor variations do not meaningfully alter the overall interpretation, demonstrating that morning insulin priming enhances PM NHGU by increasing the liver’s responsiveness to both glucose-mediated (GE/the PGS) and insulin-mediated (IA) mechanisms in the afternoon.

In exploring the cellular mechanisms underlying morning insulin priming of hepatic glucose metabolism, the rapid enhancement of NHGU within the first 30 minutes of the afternoon clamp indicates that these effects were already established at the time of secondary glucose exposure rather than developing progressively during the clamp. At the end of the PM clamp, primed animals exhibited elevated hepatic *GCK* mRNA, while total GCK protein, maximal GCK enzymatic activity, and the phosphorylation states of glycogen synthase and phosphorylase were similar between groups. The increased *GCK* mRNA is consistent with insulin-dependent transcription initiated in the morning and supports the concept that prior insulin exposure establishes a molecular program poised to enhance glucose phosphorylation capacity during subsequent hyperglycemia (38–40).

Prior work from our group demonstrated that a physiological rise in morning insulin results in increased hepatic *GCK* mRNA and GCK protein at the 330 min time point before the onset of the PM clamp (31). In the present study, the absence of elevated GCK protein at the terminal time point (480 min), despite a clear enhancement of NHGU in the Prime group (observed between 360-480 min), was unexpected. This does not preclude a central role for glucokinase, the *in vivo* activity of which is dynamically regulated by subcellular compartmentalization and protein-protein interactions. These include binding to glucokinase regulatory protein (GKRP) in the nucleus (41), feedback mechanisms involving fructose-1-phosphate (which promotes dissociation of GCK from GKRP) and fructose-6-phosphate (which promotes association of GCK and GKRP) (42), as well as the bifunctional enzyme phosphofructokinase 2/fructose-2,6-bisphosphate (43) and the pro-apoptotic mitochondrial protein BAD (44). Through these interactions and others, GCK can be sequestered or released in response to glucose availability, thereby modulating its cytosolic activity depending on the specific physiological context (45). Importantly, these complex regulatory mechanisms are not captured by terminal protein measurements or *ex vivo* activity assays. We previously attempted to assess GCK/GKRP co-localization in chow- and high-fat high-fructose-fed dogs using immunohistochemistry and fluorescence imaging, but results were not quantifiable, likely due to species-specific antigen differences. Future studies will require canine-specific protocols to isolate nuclear and cytosolic fractions from metabolically preserved tissue to accurately measure GCK and GKRP localization. The rapid, reversible GCK/GKRP interaction is highly sensitive to the metabolic milieu and difficult to preserve during tissue processing. Although freeze-clamping maintains metabolite levels, it is not optimal for fully capturing native protein-protein interactions in the dog. It is plausible that prior insulin exposure may sensitize these regulatory processes, representing potential hepatic “pre-conditioning” that could enhance glycolytic flux, an important avenue for future investigation

While we observed differences in net glycogen flux and terminal hepatic glycogen content, direct hepatic glycogen synthesis did not differ significantly, indicating that AM insulin produced only modest effects on glycogen-regulating enzyme phosphorylation at the tissue collection time. Glycogen itself may provide an additional layer of regulation, acting as a scaffold that organizes metabolic enzymes within the cell (46; 47). Subtle shifts in the localization and glycogen-binding dynamics of glycogen synthase and phosphorylase, influenced by insulin and intracellular glucose-6-phosphate (G6P), could fine-tune activity within these microdomains (48). Supporting this, fasted-to-refed rat liver demonstrates G6P-dependent translocation of glycogen synthase to glycogen clusters, independent of phosphorylation alone (49). It will be informative in future studies to measure hepatic G6P levels to assess its potential contribution to the (in)activation and subcellular redistribution of glycogen-regulating enzymes and glucokinase. Additionally, primed animals also showed higher hepatic expression of the insulin-responsive lipogenic transcription factor *SREBP-1c* (50) and its downstream target, *FAS*, at the end of the PM hyperglycemic clamp. This occurred despite only a transient rise in morning insulin, followed later by afternoon hyperglycemia, rather than the simultaneous high insulin and glucose previously thought necessary for hepatic activation of lipogenic pathways (50). These data suggest that, in addition to priming glucose-handling pathways, early insulin exposure may also sensitize hepatic lipid metabolism. Circulating lipids did not reflect the hepatic transcriptional changes, as arterial NEFA concentrations were higher in the No Prime group, yet net hepatic NEFA uptake and afternoon lipolysis were similar between groups. Future studies will be needed to define how insulin primes these lipid pathways and how this interacts with carbohydrate metabolism.

Although the precise cellular mechanism underlying morning insulin priming remains to be fully defined, GCK remains the most plausible molecular mediator. The enhancement of PM NHGU in primed animals is rapid and sustained, yet as previously mentioned, total GCK protein measured at the terminal time point (480 min) is similar between groups. This observation is consistent with the limitation of a single end-point measurement, as GCK activity *in vivo* is dynamically regulated and not fully captured by total protein abundance (51). GCK is uniquely positioned to mediate the second meal effect, as it catalyzes the rate-limiting step of hepatic glucose uptake under hyperglycemic conditions and integrates insulin-dependent transcription, a long protein half-life, and rapid activation by glucose (39; 52). Other hepatic proteins are less likely contributors, as they are not simultaneously insulin-inducible, long-lived, and acutely responsive to glucose later in the day (52). Supporting this, untargeted bulk-RNA transcriptomics from canine liver samples collected prior to the afternoon clamp indicate that GCK is among the most highly expressed transcripts following a morning insulin prime (35). While the cellular mechanisms underlying this effect remain to be fully elucidated, our flux data provides strong integrated physiological evidence for a sustained hepatic response to prior insulin exposure. Ongoing transcriptomic and LC-MS-based proteomic analyses, applied in tandem in the dog for the first time to our knowledge, are expected to further delineate the pathways involved and identify additional mediators.

An important next step is to determine how morning insulin priming is altered under conditions of metabolic dysfunction, such as in our insulin-resistant canine model induced by consumption of a high-fat, high-fructose diet for 4 weeks (7; 53), and whether these changes help explain the context-dependent expression of the second-meal phenomenon in type 2 diabetes (3; 34). Our prior work indicates that this priming effect is mediated by direct hepatic insulin action, suggesting that subcutaneous insulin delivery may not effectively engage this pathway (31). While a sufficiently large peripheral insulin dose could theoretically induce a hepatic priming effect, this would be accompanied by the risks associated with arterial hyperinsulinemia (54). Emerging approaches, including hepatopreferential insulin analogs or strategies that enhance endogenous insulin secretion, may better recapitulate this physiology (55–57), although this will require direct testing to determine whether hepatic responsiveness can be improved without inducing extensive peripheral hyperinsulinemia (54). Further work is also needed to define how nutrients (glucose, amino acids, and lipids) and hormones (insulin and glucagon) interact to coordinate hepatic metabolic memory and nutrient handling.

In summary, these studies demonstrate that a physiological rise in morning insulin primes the liver to respond more effectively to a subsequent afternoon glucose load, even in the absence of a rise in afternoon insulin. This effect is mediated in part by enhanced responsiveness to glucose-mediated signals, with additional contributions from augmented insulin action under conditions of elevated afternoon insulin. Together, these findings indicate that morning insulin exposure proportionally enhances both glucose- and insulin-mediated pathways governing NHGU. While the underlying cellular mechanisms remain to be fully defined, glucokinase represents a plausible mediator given its regulation by insulin and central role in hepatic glucose phosphorylation. These results define a hepatic mechanism by which prior insulin exposure shapes subsequent glucose handling and provide a framework for understanding insulin-induced hepatic metabolic memory. Further studies are needed to determine how this mechanism is altered in insulin-resistant states and to evaluate its relevance in humans.

## Author Contributions

H.W., D.S.E., and A.D.C. participated in the design of experiments; H.W. directed all experiments and collected data; B.F., K.Y., and T.H. participated in the experiments; B.F. and K.Y. were responsible for surgical preparation and oversight of animal care; H.W. and M.S.S. carried out biochemical and tissue analysis; H.W., D.S.E., and A.D.C. interpreted experimental results; H.W. prepared figures and drafted the manuscript; H.W., D.S.E., G.K., and A.D.C. edited and revised the manuscript; All authors approved the final version of the manuscript.

## Guarantor Statement

A.D.C. is the guarantor of this work and, as such, has full access to all the data in the study and takes responsibility for the integrity of the data and the accuracy of the data analysis.

## Duality of Interest

A.D.C. has research contracts with Alnylam and Abvance, as well as grants from the NIH. A.D.C. is a consultant to Novo Nordisk, Fractyl, Sensulin, and Thetis. No other individuals have conflicts of interest relevant to this article.

## Funding

This work was supported by National Institutes of Health (NIH) grant R01DK131082. H.W. was supported by NIH grant F31DK142297 and by the Vanderbilt University Training Program in Molecular Endocrinology NIH grant T32DK007563. Hormone analysis was completed by the Vanderbilt University Medical Center Analytical Services Core, supported by the NIH Diabetes Research Center grant DK020593. Vanderbilt’s Large Animal Core provided surgical expertise, supported by the NIH Grant DK020593.

## Prior Presentation

This work was orally presented at the American Diabetes Association 2025 meeting. The abstract can be viewed at 10.2337/db25-362-OR. The graphical abstract was created with BioRender and published with permission.

---

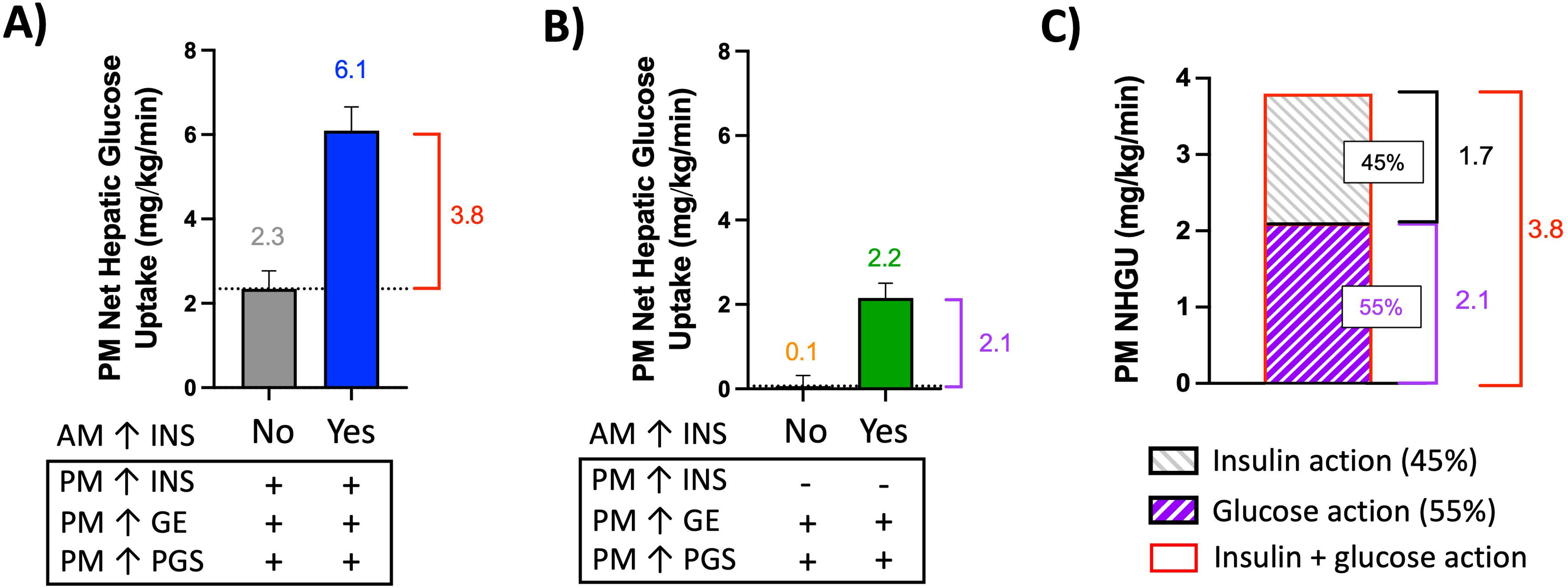

## Notes

https://figshare.com/articles/journal_contribution/Supplemental_Materials_and_Supporting_Data_Values_File_for_b_Morning_Elevation_in_Insulin_Enhances_Afternoon_Postprandial_Glucose_Action_and_Insulin_Action_in_Canines_b_/31049578

### Summary of Updates

Figures were updates to fully reflect the work and the discussion was expanded.

