## Supplemental for "Morning Elevation in Insulin Enhances Afternoon Hepatic Glucose Disposal in Dogs by Increasing Both Insulin Signaling and Glucose Action"

**Hepatic glycogen content**

Liver glycogen content was measured using a modification of the method of Keppler and Decker (1; 2). Frozen liver tissue (~180 mg) was weighed under liquid nitrogen to prevent thawing and enzymatic degradation. Tissue weight (mg) was used to calculate the volume of 0.6 N perchloric acid (PCA) required for homogenization (volume = tissue weight × 5). Samples were homogenized on ice in the calculated volume of PCA. From each homogenate, 200 µL aliquots were transferred into separate tubes and neutralized with 100 µL of potassium bicarbonate. To quantify glycogen, 500 µL of amyloglucosidase solution (2 mg/mL in 0.4 M sodium acetate buffer) was added to experimental aliquots and incubated at 40°C in a shaking water bath for 2 hours to hydrolyze glycogen to glucose. Duplicate control samples lacking amyloglucosidase were processed in parallel to account for free glucose in the homogenate. Glycogen content was calculated as glucose released in enzyme-treated samples minus glucose in paired control samples.

Standards were prepared using oyster glycogen (4, 5.5, and 7 mg) dissolved in 1 mL of 0.6 N PCA and processed alongside samples. After incubation, samples were cooled on ice and centrifuged at 3,000 rpm for 5 minutes. For samples containing tracer, 500 µL of supernatant was transferred to scintillation vials for [³H]-glycogen counting using the same evaporation, reconstitution, and scintillation methods applied to plasma 3-[³H]-glucose. Remaining aliquots were analyzed for glucose concentration using an Analox GM9 glucose analyzer. All solutions (PCA, potassium bicarbonate, amyloglucosidase) were prepared fresh on the day of assay.

**RNA extraction, cDNA synthesis, and quantitative real-time PCR**

Total RNA was isolated from canine liver tissue to assess hepatic gene expression. Approximately 50 mg of frozen liver tissue was homogenized in 1 mL of Tri-reagent (Sigma-Aldrich, St. Louis, MO) according to the manufacturer’s instructions. RNA purification was performed using the Direct-zol RNA Miniprep Kit (Zymo Research, Irvine, CA) with final elution in 35 µL of nuclease-free TE buffer. RNA yield and purity were determined by spectrophotometry, with A260/A280 ratios >1.8 considered acceptable. RNA integrity was verified by visualization on ethidium bromide–stained agarose gels. All procedures were conducted using RNase/DNase-free reagents and consumables. First-strand cDNA was synthesized from 1 µg of total RNA using the High-Capacity cDNA Reverse Transcription Kit (Applied Biosystems, Foster City, CA) following the manufacturer’s instructions. cDNA was stored at –80°C until use. Primers for target and reference genes were designed with Beacon Designer software (Premier Biosoft, Palo Alto, CA), verified for specificity with BLAST, and further confirmed by melt curve analysis. Primer efficiency was within the range of 91%–96%.

Quantitative real-time PCR was carried out on a CFX96 Real-Time PCR Detection System (Bio-Rad, Hercules, CA) using SsoAdvanced Universal SYBR Green Supermix (Bio-Rad). Each 25 µL reaction contained 100 ng cDNA template, 12.5 µL SYBR Green Supermix, 0.4 µM forward and reverse primers, and nuclease-free water. Thermal cycling conditions were as follows: 95°C for 3 minutes, followed by 39 cycles of 95°C for 10 seconds and 55°C for 30 seconds. All reactions were run in duplicate. Melt curve analysis was performed to ensure amplification specificity. Relative gene expression was calculated using the 2^–ΔΔCt method, with GAPDH as the reference gene (3). Data represent the mean of 2–3 independent PCR runs. Tissue from the left central and left lateral lobes was analyzed, as these lobes account for ~50% of total liver mass and provide a representative assessment of hepatic gene expression. Primer details can be viewed in Table 1 below.

**Supplemental Table 1 - Primer details for quantitative PCR of selected genes**

| **Gene Name** | **Forward Primer (5’ 🡪 3’)** | **Forward Melting Temperature (Tm)** | **Reverse Primer (5’ 🡪 3’)** | **Reverse Melting Temperature (Tm)** |
| --- | --- | --- | --- | --- |
| *GCK* | CAGAGGGGACTTTGAAATG | 59.6°C | CTGCATCTCCTCCATGTAG | 58.3°C |
| *PCK1* | AGCTTTCAATGCCCGATTTCCAGG | 73.1°C | TCAGCTCGATGCCGATCTTTGACA | 73.7°C |
| *G6PC1* | CCTTTATTCCTCTTTC | 50.7°C | GGTGTTGCTATAGTAGTC | 46.0°C |
| *SREBP-1c* | GTGAAGGCAGCGGGTATCAG | 66.8°C | TCTCAGTGTCCACTACCAGAGG | 63.2°C |
| *FAS* | TACTGGAGGGGCCAGTGCATCA | 72.9°C | GTCCCGAGATGGTCACTGTGTC | 68.1°C |
| *GAPDH* | TGTCCCCACCCCCAATGTATC | 69.8°C | CTCCGATGCCTGCTTCACTACCTT | 70.2°C |

**Western blotting**

Frozen liver tissue (~100 mg) was homogenized in 1 mL of ice-cold homogenization buffer (20 mM Tris, 200 mM NaCl, 50 mM NaF, 1 mM EDTA, 1 mM EGTA, 10% glycerol, 1% SDS, pH 7.2) supplemented with protease and phosphatase inhibitors (Sigma-Aldrich, St. Louis, MO). Homogenates were centrifuged at 3,000 × g for 10 minutes at 4°C, and supernatants were collected. Protein concentrations were determined using a BCA protein assay (Bio-Rad, Hercules, CA). Protein samples were adjusted to 4 µg/µL in Laemmli buffer, heat-denatured at 95°C for 5 minutes, and stored at –80°C. Equal amounts of protein (20 µg per lane) were separated on 4-12% Criterion TGX gels (Bio-Rad) and transferred to nitrocellulose membranes using a semi-dry Trans-Blot SD transfer system (Bio-Rad) in Towbin buffer (25 mM Tris, 192 mM glycine, 20% methanol, pH 8.3) for 25 minutes at 15 V. Membranes were stained with Ponceau S solution to visualize total protein, imaged with a ChemiDoc system (Bio-Rad), and normalized to Ponceau S staining intensity. Membranes were then washed with TBST (10 mM Tris, 150 mM NaCl, 0.2% Tween-20, pH 7.5) and blocked for 1 hour at room temperature with 5% (w/v) bovine serum albumin in TBST. Blots were incubated overnight at 4°C with primary antibodies diluted in blocking buffer. The following primary antibodies were used in western blotting analysis: pAkt (Ser473, #9271, Cell Signaling Technology, Danvers, MA), tAkt (#9272, Cell Signaling Technology), pGS (Ser641, #3891, Cell Signaling Technology), tGS (#98348, Cell Signaling Technology), pGP (Ser15, #ab227043, Abcam, Cambridge, UK), tGP (#ab198268, Abcam), and GCK (#sc-17819, Santa Cruz Biotechnology). Primary antibody dilutions were all 1:5,000, except for GCK (1:10,000). After three washes in TBST (x5 min), membranes were incubated with 1:5,000 Anti-rabbit HRP-conjugated secondary antibodies (Promega, Madison, WI) for 1 hour at room temperature, followed by three additional washes (x5 min) in TBST. Proteins were detected using ECL Plus reagents (GE Healthcare, Piscataway, NJ) and imaged with a ChemiDoc system. Band intensities were quantified with ImageJ software (NIH, Bethesda, MD), normalized to total protein detected by Ponceau S. Data represent the mean of two blots per protein. Western blotting was performed on tissue from the left central and left lateral lobes, which together account for ~50% of total liver mass in the dog.

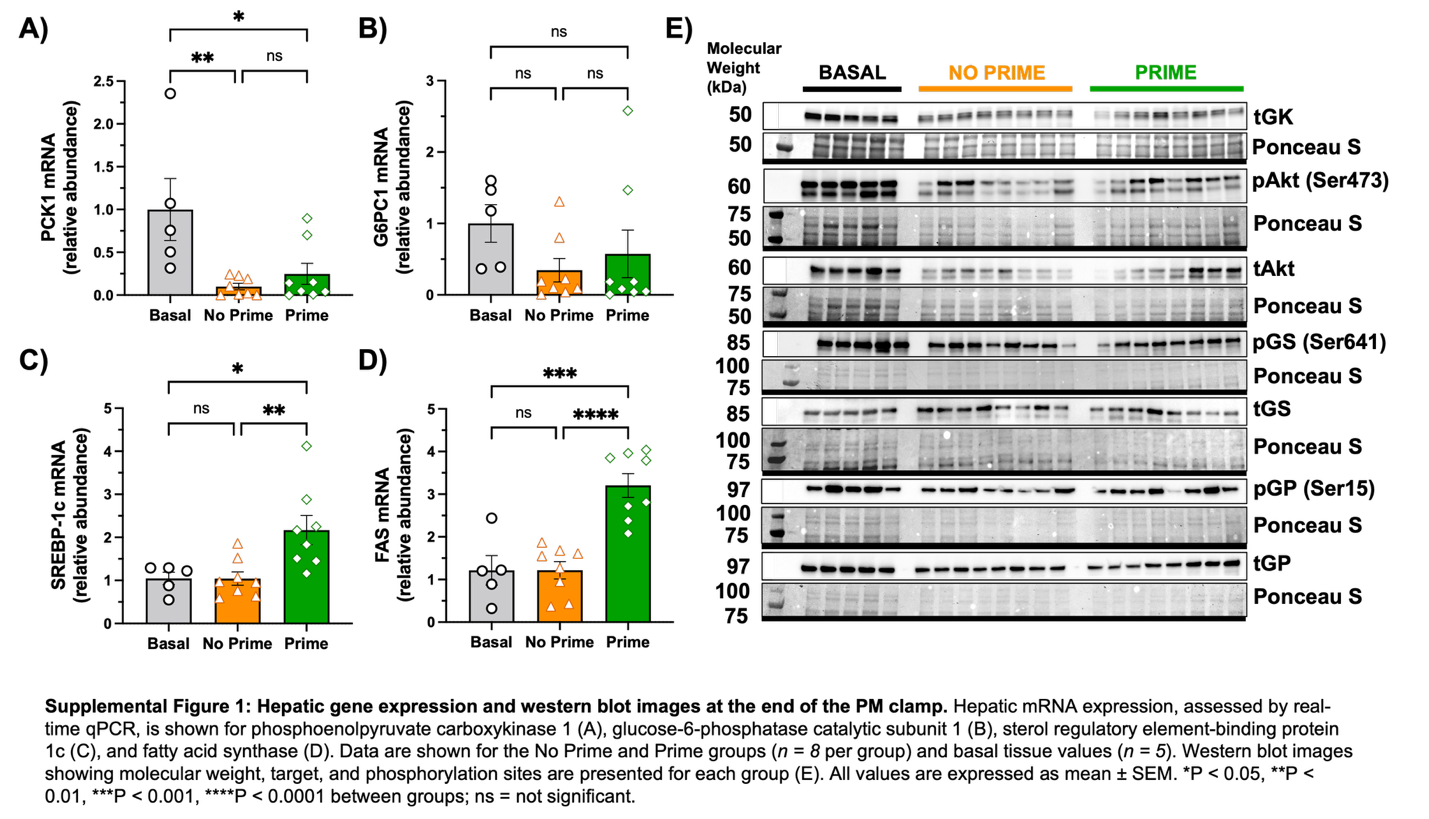

**Glucokinase Activity Assay**

Glucokinase activity was measured using an assay adapted from Shiota et al. (4). 60 mg of frozen canine liver tissue was homogenized in 1.5 mL of homogenization buffer containing 50 mM HEPES, 100 mM KCl, 5 mM MgCl_2_, 1 mM EDTA, and 2.5 mM dithiothreitol. Homogenates were centrifuged at 30,000 rpm for 45 minutes at 4 °C using a TLA-55 rotor, and the resulting supernatant was collected and maintained on ice until analysis. For the assay, 10 µL of supernatant was added to 190 µL of reaction medium (37 °C; 2% albumin, 50 mM HEPES, 100 mM KCl, 7.5 mM MgCl_2_, 2.5 mM DTT, 0.5 mM NAD⁺, 5 mM ATP, and 2 units of glucose-6-phosphate dehydrogenase). Reactions were performed in the presence of varying glucose concentrations (0, 0.5, 7, and 100 mM). Following substrate addition, plates were incubated at 37 °C, and absorbance was recorded at 340 nm in 5-minute intervals over a 30-minute period. GCK activity was calculated at both ½ Vmax and Vmax. Baseline liver samples from five dogs that were fasted overnight were included for comparison.
